# Mechanisms of transcription control by distal enhancers from high-resolution single-gene imaging

**DOI:** 10.1101/2023.03.19.533190

**Authors:** Lingling Cheng, Chayan De, Jieru Li, Alexandros Pertsinidis

## Abstract

How distal enhancers physically control promoters over large genomic distances, to enable cell-type specific gene expression, remains obscure. Using single-gene super-resolution imaging and acute targeted perturbations, we define physical parameters of enhancer-promoter communication and elucidate processes that underlie target gene activation. Productive enhancer-promoter encounters happen at 3D distances δ200 nm - a spatial scale corresponding to unexpected enhancer-associated clusters of general transcription factor (GTF) components of the Pol II machinery. Distal activation is achieved by increasing transcriptional bursting frequency, a process facilitated by embedding a promoter into such GTF clusters and by accelerating an underlying multi-step cascade comprising early phases in the Pol II transcription cycle. These findings help clarify molecular/biochemical signals involved in long-range activation and their means of transmission from enhancer to promoter.

## Main Text

Activation of transcription by distal enhancers uncouples *cis*-regulatory information from the promoter-proximal region and enables controlling the same gene in intricate spatio-temporal patterns of cell- and tissue-specific expression (*1, 2*). Major progress has been made in describing genome structure-function relations and the interconnectedness of 3D genome topology and long-range gene regulation (*3-8*). Understanding how enhancer-promoter communication is physically achieved in 3D nuclear space remains a key problem(*9*). Recent advances in single-molecule and single-gene imaging methods have revealed a striking nano-scale organization of focally accumulated enhancer-associated regulatory factors (RFs), within 100-200 nm of active genes (*10, 11*). The same studies also revealed one-to-one correspondence between clustering of certain RFs and the frequency of transcriptional bursting(*10*), as well as coordinated kinetics of two linked promoters that share a partially overlapping pool of clustered RFs(*11*). These results had indicated that a local nano-scale environment(*12*) might contain activating signals that facilitate transcriptional bursting of embedded promoters. However, how transcription activity relates to exact distances between promoter and nanoscale RF clusters has thus far not been quantified. The previous imaging studies did not address how transcriptional bursting kinetics are quantitatively affected when promoter-enhancer spatial relationships are modulated and what defines the dynamic range of productive promoter-enhancer encounters.

On a fundamental level, enhancers are thought to activate transcription by regulating processes that facilitate the function of the RNA Polymerase II machinery at target promoters (*13*). Consistent with this notion, enhancers control the frequency of transcriptional bursts (*14-16*), regulating how often a promoter is active (*17*). However, the precise molecular transactions and/or biochemical activities that are facilitated by enhancer-promoter communication, as well as the molecular and biophysical means for achieving such facilitation have not been well characterized. Understanding the nature of activating signals transmitted from enhancer and associated RFs to promoter and the Pol II machinery, to initiate transcription bursts, is thus of utmost importance. To address these questions, we sought to directly visualize at high resolution key aspects of transcription activation by distal enhancers: promoter-enhancer distances, nano-scale organization of regulatory factors and Pol II machinery components, and real-time kinetics of nascent RNA production. We further combine real-time imaging with systematic acute perturbations using targeted protein degradation and small-molecule inhibitors. This approach unveils key quantitative and mechanistic insights into promoter-enhancer interactions, transcription kinetics and the underlying molecular and/or biochemical processes involved.

### Promoter-enhancer spatial encounters modulate on-off bursting kinetics

To establish experimental systems where spatial relations between promoter and enhancer can be modulated, we turn our attention to cohesin, a genome architectural factor(*18*) that can also act as facilitator of promoter-enhancer interactions (*19-22*). We establish knock-in mouse embryonic stem cells (mESCs) where the endogenous Rad21 cohesin sub-unit can be rapidly degraded (Fig. 1**A-B**, Supplementary Fig. 1**A**) using the dTAG targeted protein degradation system (*23*). Simultaneously, we use live-cell real-time readout of nascent transcription activity via the MS2-MCP system (*24*) to evaluate the effects of these perturbations on transcription bursting (Fig. 1**A**).

**Figure 1.**
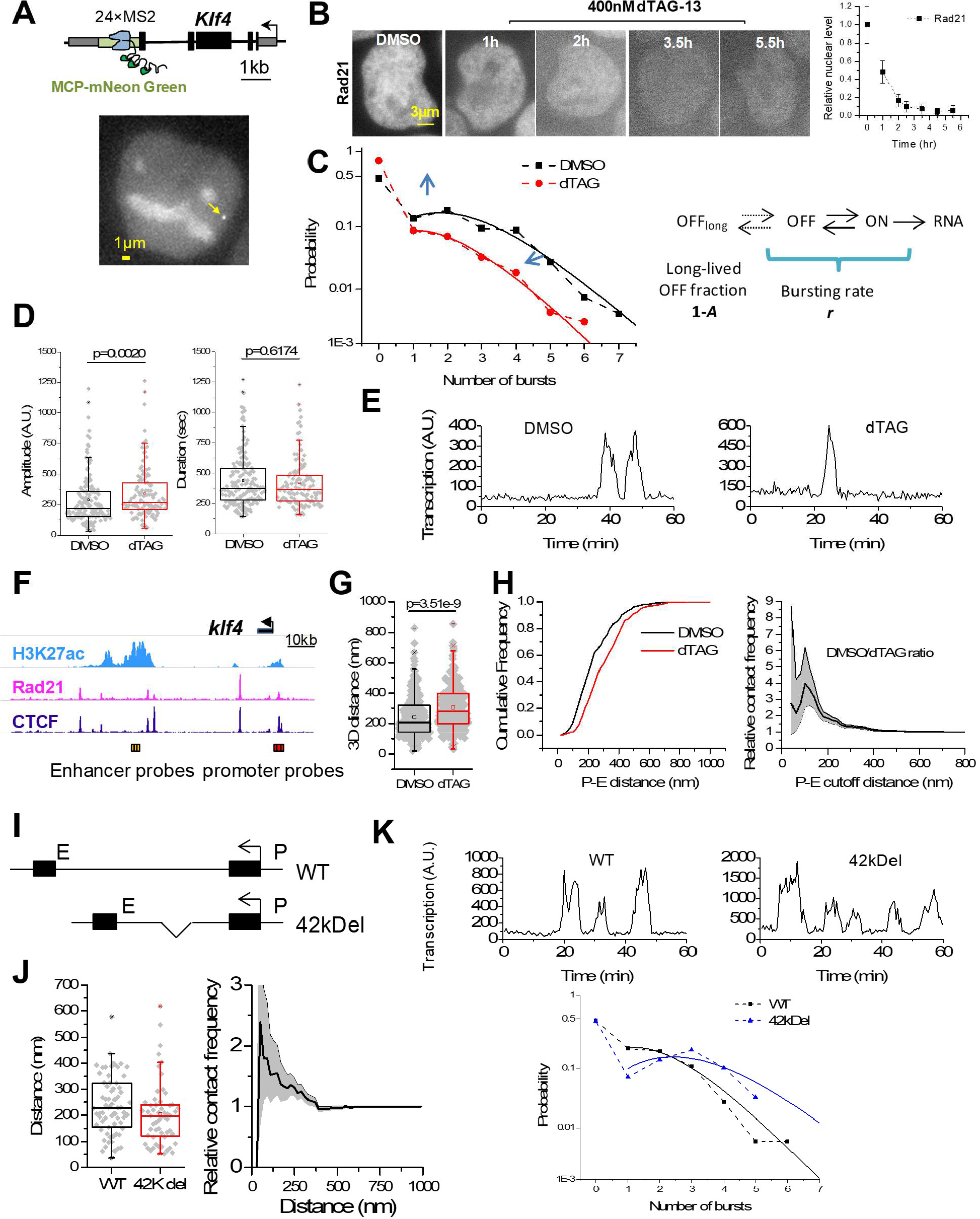
Promoter-enhancer spatial encounters modulate on-off bursting kinetics. (**A**) Schematic of the *Klf4* locus integrated with 24×MS2 (green rectangle) and representative image of a nucleus with an actively transcribed locus. Black rectangles are exons and gray rectangles are 5′ and 3′ UTR. The stem loops formed by MS2 RNA can be recognized by MCP-mNeonGreen. The yellow arrow points to *Klf4* nascent transcription site. (**B**) Rad21-Halo staining showing the time-course of treatment with dTAG-13 (400nM). Control cells are treated with DMSO for 6 hrs. Scale bar = 3µm. The left panel shows quantification of relative Rad21 nuclear levels, normalized to DMSO. (**C**) Distribution of *Klf4* bursts in DMSO-(*n*=119 traces from 303 cells) and dTAG-treated (*n*=110 traces from 780 cells) Rad21-FKBP^F36V^-halo KMG cells. Each data point represents the probability that a certain number of bursts, *n*_bursts_, was observed in 1 hr. Solid lines indicate fits to a Poisson distribution for the data with *n*_bursts_≥1. The resulting two parameters (A, *r*) are: DMSO (0.6, 2.29), dTAG (0.28, 1.59). (**D**) Box plots of burst amplitude and duration. Each gray dot represents one burst. Data points are from 3 independent experiments with total *n*= 160 and 117 bursts for DMSO and dTAG, respectively. p-values are calculated based on a Wilcoxon rank-sum test. (**E**) Two intensity traces of transcription sites from DMSO- and dTAG-treated Rad21-FKBP^F36V^-Halo KMG cells. (**F**) Profiles of H3K27ac, Rad21 and CTCF ChIPseq signals at the *Klf4* Locus (published datasets GSM1526287, GSM2418859 and GSM2418860, respectively.). The red and yellow rectangles indicate the positions of DNA FISH probes (tagging promoter and -55kb enhancer regions, respectively). Each set of probes spans around 3kb. (**G**) 3D promoter-enhancer distances based on OligoPAINT FISH, for DMSO vs. dTAG treated KMG cells (mean ± SD: WT 240±132, dTAG 305 ± 140). *n*= 280 and 257, respectively. (**H**) Left: Cumulative distribution of enhancer– promoter 3D distances in Rad21-FKBP^F36V^-halo KMG cells at the *Klf4* loci. Right: relative contact frequency distribution estimated from the DMSO/dTAG ratio of cumulative distance distributions. Grey shaded area indicates binomial 95% confidence intervals. (**I**) Schematic of 42 kb truncation between *Klf4* promoter and +55 kb downstream enhancer (42kDel). (**J**) Left: 3D enhancer-promoter distances (mean ± SD: WT 237±111, 42kDel 205 ± 121). Right: relative contact frequency, estimated by the ratio 42kDel/WT of cumulative distance distributions. Grey shaded area indicates binomial 95% confidence intervals. (**K**) (Top): Two intensity traces of transcription sites from WT and 42kDel KMG cells. (Bottom): Probability of observing certain number of bursts in an 1 hr observation time, in WT (*n*=93 traces from 85 cells) and 42kDel (*n*=110 traces from 104 cells) KMG cells. Solid lines: fits to Poisson distribution, for the data with *n*_bursts_≥1.

Detailed analysis of transcription bursting kinetics of selected pluripotency genes in mESCs reveals deviations from the popular two-state random telegraph model (*25, 26*). Rather, bursting kinetics can be better described with at least one additional long-lived off state (*27, 28*), resulting in two distinct sub-populations: a sub-population that is exhibiting on-off bursting with largely a fixed frequency (characterized by Poisson-distributed bursts in a given time interval), and a sub-population that is effectively silent during the observation time (1 hr) (Fig. 1**C**, Supplementary Fig. 1**B-C**).

Rapid depletion of Rad21 results in ∼2-fold decrease of the bursting frequency *r*, and ∼2-fold increase of the long-lived off-state sub-population fraction, *A*, for bursting of *Klf4* (Fig. 1**C**). At the same time, burst amplitude is only very modestly affected (∼10%) while burst duration remains unaffected (Fig. 1**D**). Similar behavior is observed for *Sox2* and *Tbx3* (Supplementary Fig. 1**B-C**). These results indicate that cohesin controls transcription activity in an almost strictly digital on-off manner – by increasing burst initiation rate and by preventing entry into long-lived off states.

To investigate the underlying changes in the spatial relations between promoter and enhancer that are associated with the reduced transcription activity upon Rad21 loss, we use multi-color super-resolution imaging with OligoPAINT FISH probes (*29*) (Fig. 1**F**, Supplementary Fig. 1**D-F**). Nanometer multi-color 3D distance measurements between probes tagging the *Klf4* promoter and the 55kb down-stream *Klf4* enhancer indicate a unimodal distribution of promoter-enhancer distances, with a median of 207 nm (Fig. 1**G**). Rad21 depletion results in a modest shift of the overall distribution towards larger distances, with the median increasing to 279 nm (Fig. 1**G**).

To better understand the changes in promoter-enhancer spatial proximity conferred by loss of Rad21, we quantify the relative cumulative distance distribution D_*r*_(*r*), obtained as the ratio of the cumulative distance distributions for control cells treated with DMSO vs. cells treated with dTAG. This ratio effectively reports the relative abundance of distances below a specific distance threshold, *r*. D_*r*_(*r*) is ∼1 for threshold distances larger than ∼200 nm, and gradually increases above 1 as the distance threshold decreases (Fig. 1**H**). This analysis reveals that cohesin-mediated processes increase encounters that bring promoter and enhancer to <200 nm from each other. Together with the observed transcriptional responses, these results suggest that these close-range encounters are important for transcription activation.

Since decreased frequency of enhancer-promoter encounters upon Rad21 loss leads to attenuated on-off bursting kinetics, we reason that increased spatial proximity will lead to more frequent bursting. To test this idea, we created genome-edited mESCs where 42kb of downstream DNA between *Klf4* and its +55kb enhancer have been deleted (42kDel) (Fig. 1**I**). In 42kDel mESCs we observe significantly reduced physical promoter-enhancer distance compared to WT (Fig. 1**J**). The relative cumulative distance distribution D_*r*_(*r*) of 42kDel vs. WT indicates increased frequency of encounters that bring promoter and enhancer to <200 nm (Fig 1**J**). At the same time, 42kDel cells show increased bursting frequency (Fig 1**K**). Taken together with the results from Rad21 depletion, our observations suggest that modulation of enhancer-promoter spatial encounters controls the on-off transcriptional bursting kinetics.

### A multi-step cascade underlies the initiation of transcriptional bursts

The more detailed understanding of how cohesin depletion and decreased promoter-enhancer interactions affect nascent transcription kinetics paves the way for further elucidating molecular mechanisms by which enhancers activate transcription. Specifically, we reason that promoter-enhancer interactions likely facilitate molecular processes that are involved in the initiation of transcriptional bursts. Disturbing processes that are facilitated by enhancer-promoter communication would then resemble the effects of cohesin loss on the on-off digital control of nascent transcription activity.

Molecular processes often envisioned to be facilitated by promoter-enhancer interactions include deposition of transcription factors, epigenetic marks, or remodelling of nucleosomes to increase DNA accessibility (*13, 30*). Once these processes have been completed, the promoter is in an open and active state, and the transcription cycle of Pol II (*31*) can proceed efficiently and rapidly; multiple Pol II molecules are released into productive elongation in close succession, producing a burst of multiple RNA synthesis. This picture is consistent with the view that enhancers mostly control burst frequencies (i.e. the probability that the promoter is active), while promoters mostly control burst sizes (i.e. the amount of RNA that is produced while the promoter is active) (*14, 15*).

Based on these proposed roles of enhancers and promoters in controlling bursting kinetics, we predict that perturbations of enhancer-associated transcription factors and chromatin regulators would mostly affect burst frequency, while disrupting the general Pol II transcription machinery would mostly affect burst amplitude and/or duration. To test these ideas, we perform a series of systematic perturbations using targeted protein degradation and small-molecule chemical inhibitors (Fig 2**A-D**, Supplementary Fig. 2, Supplementary Fig. 3). We focus on multiple different components spanning a large part of the transcription process and its regulation: from sequence-specific DNA binding transcription factors (Sox2), to chromatin regulators (BET proteins/Brd4), to components of the Pre-initiation complex (Tbp/Trf2, TFIID, Mediator, Pol II, TFIIF, TFIIH), to factors controlling the release of Pol II into productive elongation (pTEFb/Cdk9).

**Figure 2.**
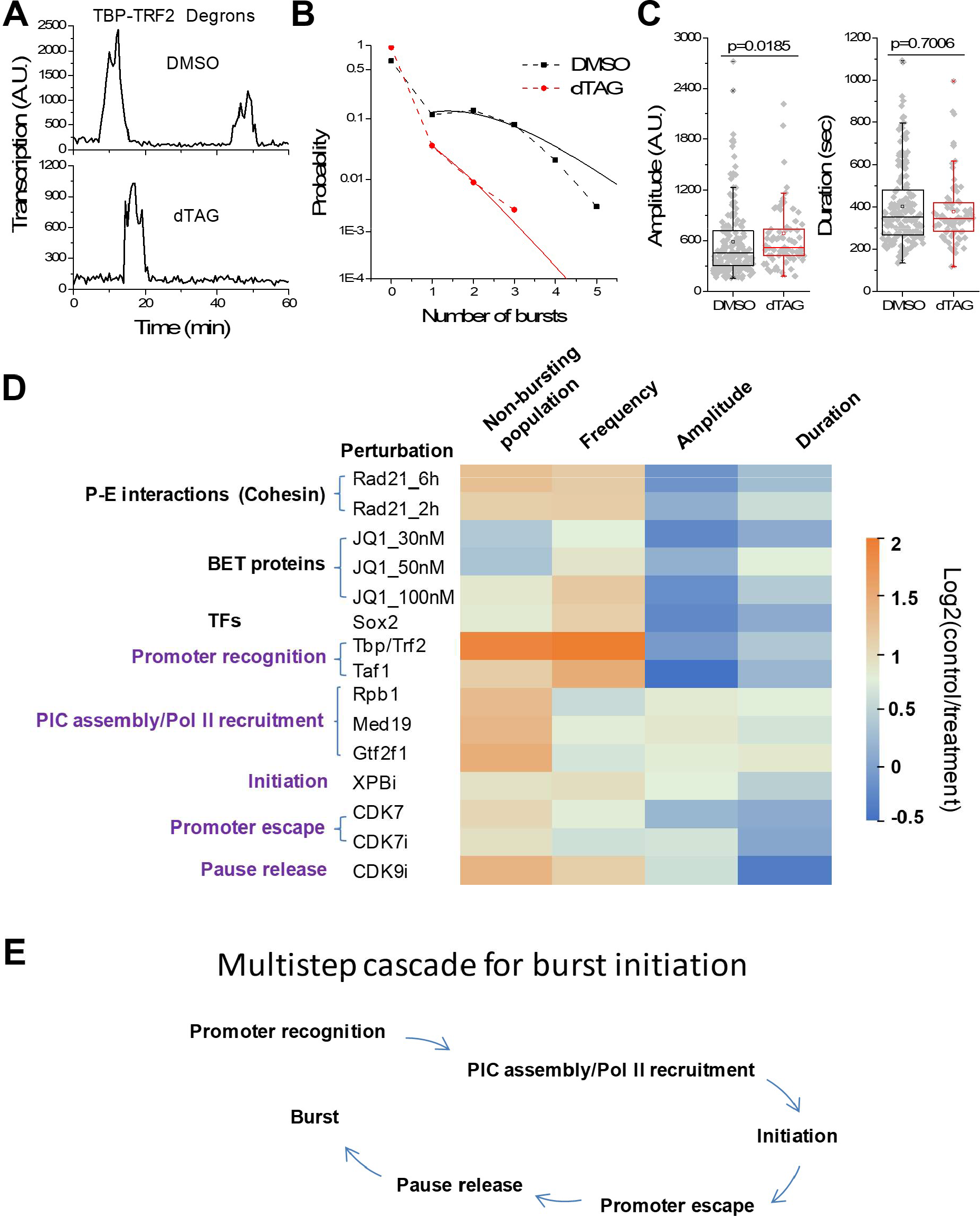
A multi-step cascade underlies the initiation of transcriptional bursts. (**A**) Two intensity traces of transcription sites from DMSO- and dTAG-treated Tbp-Trf2-FKBP^F36V^ KMG cells. (**B**) Probability of observing certain number of bursts in an 1 hr observation time, in DMSO-(*n*=114 traces from 161 cells) and dTAG-treated (*n*=72 traces from 738 cells) Tbp-Trf2-FKBP^F36V^ KMG cells. Solid lines: fits to Poisson distribution, for the data with *n*_bursts_≥1. The resulting two parameters (A, *r*) are: DMSO (0.44 and 1.91), dTAG (0.12 and 0.49). (**C**) Box plots of burst amplitude and duration. Each gray dot represents one burst. Data points are from two independent experiments with total *n*= 175 and 73 bursts for DMSO and dTAG, respectively. p-values are calculated based on a Wilcoxon rank-sum test. (**D**) Heat-map of fold-change of bursting parameters (non-bursting population, frequency, amplitude and duration) from different perturbations in KMG cells, including dTAG induced degradation (Rad21, Sox2, Tbp-Trf2, Taf1, Med19, Gtf2f1 and CDK7), Halo-PROTAC-E induced partial degradation (40nM) of Rpb1, small amount of inhibitors [100nM THZ1 (CDK7i) and triptolide (XPBi), 5nM NVP-2(CDK9i)], and JQ1. Data are normalized by Log2 (control/treatment). The data include 2-3 independent experiments. (**E**) Multi-step burst initiation cascade schematic, including the early phases of the Pol II transcription cycle.

Disruption of enhancer-associated regulatory factors markedly affects the on-off behavior of transcriptional bursts: targeted degradation of Sox2 and chemical inhibition of BET proteins/Brd4 decreases bursting frequency and increases the non-busting population, while leaving burst amplitude and duration mostly unaffected (Supplementary Fig. 3, Fig 2**D**). The effects are very similar to the behaviour seen after attenuation of promoter-enhancer interactions upon Rad21 loss. These results further extend previous imaging studies that linked enhancer-associated RFs to transcription bursting (*10*), and are consistent with the notion that enhancers control bursting frequency (*14-16*).

Contrary to our predictions of burst size (amplitude and/or duration) modulation, perturbations of the general Pol II transcription machinery acting at the promoter also predominantly affect on-off bursting behavior. Disrupting the first step in the Pol II transcription cycle (*31*), promoter recognition, closely resembles the effects of perturbing enhancer-promoter communication and enhancer-associated RFs. Targeted degradation of Tbp/Trf2 and the Taf1 TFIID subunit results in dramatic (2-4-fold) reduction on the frequency of transcription bursts, as well as an increase in the non-bursting population. However, when bursts are infrequently observed, burst amplitude is only modestly affected (16-40% increased), while burst duration remains unchanged (Fig 2**A-C**, Supplementary Fig. 3). These results suggest the promoter recognition is one of the processes that occur as part of the burst initiation pathway.

Disrupting subsequent steps of the Pol II cycle at the promoter also resembles effects seen by disruption of enhancer-promoter communication and enhancer-associated RFs. We perturb (i) PIC assembly and Pol II recruitment (via degradation of the Mediator subunit Med19, the TFIIF subunit Gtf2f1, and partial degradation of the Pol II subunit Rpb1); (ii) Initiation (via chemical inhibition of the ATPase of the XPB TFIIH subunit); (iii) Promoter escape (via targeted degradation or chemical inhibition of the Cdk7 TFIIH subunit); (iv) Promoter-proximal pause release (via chemical inhibition of the Cdk9 pTEFb subunit). All these perturbations affect the on-off bursting kinetics – via decreasing bursting frequency and/or increasing the non-bursting population (Fig 2**D**, Supplementary Fig. 3). At the same time, some perturbations (Rpb1, Med19, and Gtf2f1) also reduce burst amplitude (and to a lesser degree affect burst duration), possibly reflecting rate-limiting steps that occur after a burst has initiated. Overall, perturbations of the general Pol II machinery reveal that burst initiation is a cascade with multiple successive biochemical steps, and, unexpectedly, it involves many of the early steps in the Pol II transcription cycle (Fig. 2**E**).

### Increased bursting frequency is associated with proximity of target gene to Pol II GTF clusters that form at distal enhancers

Previous single-molecule nanoscopy studies described focal accumulation of several key Pol II regulatory factors at transcription sites, but did not elucidate downstream molecular processes that might link enhancer RF clustering to control of target promoter bursting. Our discovery that the activities of several Pol II GTFs are involved in the initiation of transcriptional bursts prompted us to analyze the distribution of components of the transcription machinery at active transcription sites in more detail. Strikingly, single-gene imaging reveals certain Pol II GTFs focally accumulate in the vicinity of transcription sites, akin to enhancer-associated RFs (Fig. 3**A**).

**Figure 3.**
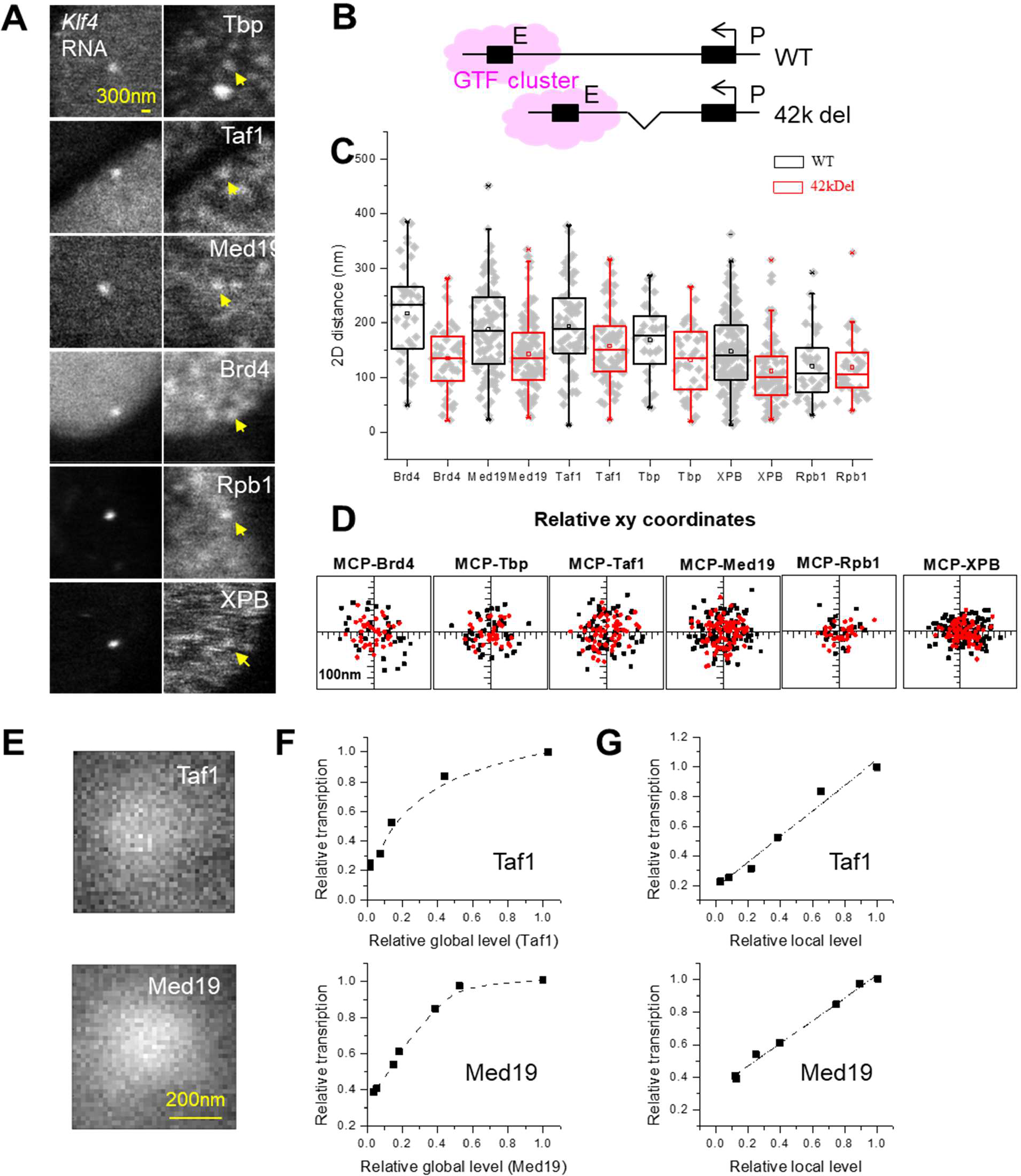
Transcription bursting is enhanced by proximity to Pol II GTF clusters that form at distal enhancers. (**A**) Live-cell single-gene imaging (3.5 μm × 3.5 μm ROIs) shows co-localized MCP-mNeonGreen and Halo-Rpb1, -Tbp, -Med19, -Taf1, -XPB and SiR -Brd4 foci (yellow arrows) at the *Klf4* locus. (**B**) Hypothesis that enhancer formed GTF cluster will be closer to the promoter in 42kDel cells. (**C**) Relative MCP-Brd4, -Taf1, -Tbp, -Med19, -XPB and -Rpb1 2D distances, (mean ± SD): Brd4 217 ± 91(135 ± 63)nm, Taf1 194 ± 83(158 ± 67)nm, Med19 188± 83(142 ± 67)nm, Tbp 169± 67(133 ± 63)nm, XPB 148± 73(111 ± 45)nm, Rpb1 121 ± 65(119 ± 58)nm, n = 37(40), 64(64), 91(93), 31(35), 133(77), 32(34), in WT(42kDel) cells, respectively. Each point corresponds to the distance of a single transcription site. All data points are from 2-3 independent experiments. (**D**) Relative *xy* coordinates corresponding to the points in (**C**). (**E**) Increased local concentration of GTFs in the vicinity of *Klf4*. Density profiles were obtained from summing multiple transcription site images aligned at the center of the *Klf4* nascent RNA; n= 68 and 106 for Taf1 and Med19, respectively, from two and three independent experiments. (**F**,**G**) Relative transcription bursting vs. global (**F**) and local (**G**) GTF concentrations, respectively. Dashed lines correspond to Hill (**F**) and linear (**G**) curve fits.

To better understand where these GTF clusters form, we analyze the WT *Klf4* gene and the 42kDel derivative, which features significantly reduced physical promoter-enhancer distance compared to WT (Fig 1**I-K**). We reason that if GTF clusters form at the distal enhancer, they will appear correspondingly closer to the MCP-tagged nascent *Klf4* RNA for the 42kDel compared to the WT construct (Fig. 3**B**). Alternatively, if GTFs cluster at the *Klf4* promoter (or in some other region uncoupled from the *Klf4* locus), the GTF-nascent RNA distances would be similar in the two constructs. As controls, we also examine RF clusters that are expected to be associated with the enhancer (*10, 11*), and Pol II clusters, previously shown to contain mostly elongating Pol II at the gene body (*10, 11*). Our measurements show that both GTF and RF clusters move closer to the transcription site (tagged via the nascent RNA) for 42kDel compared to the WT, while the Pol II clusters show same distances for both constructs (Fig. 3**C-D**). We also analyze the distribution of GTFs in a cell line where a TetO array has been integrated in the vicinity of the +100kb distal *Sox2* enhancer (*Sox2* control region; SCR). Similarly, GTF and RF clusters are better co-localized with the TetO array marking the SCR, while Pol II clusters (reflecting elongating Pol II) are better colocalized with the *Sox2* nascent RNA (Supplementary Fig. 4**A-D**). These results reveal that certain Pol II GTFs form clusters at distal enhancers. Together with the observation of more frequent transcriptional bursts in 42kDel vs. WT mESCs, our results further indicate that proximity of a promoter to GTF clusters at the enhancer is associated with increased transcription activity.

Clustering locally creates a region of high GTF concentration (Fig. 3**E**). We reason that if this high local GTF concentration controls promoter activity, it would be a better determinant of bursting kinetics compared to the global nuclear GTF concentration. To modulate the global and local GTF abundances we titrate dTAG. For the GTFs Taf1 and Med19 we achieve a range of total nuclear abundances and local cluster sizes at the *Klf4* locus (Supplementary Fig. 5**A-B**). Over this titration range, GTF cluster-*Klf4* transcription site distances remain unchanged (Supplementary Fig. 5**C**). This is consistent with an absence of Taf1 and Med19 roles in promoter-enhancer interactions, and indicates that over our experimental dynamic range, cluster size is the main parameter affecting local Taf1 and Med19 concentrations. *Klf4* transcriptional response to gradual Taf1 and Med19 loss exhibits “ buffering”, showing almost no response over >2-fold changes in global Taf1 and Med19 nuclear concentration (Fig. 3**F**). In stark contrast, *Klf4* bursting frequency is directly proportional to the local Taf1 and Med19 cluster sizes (Fig. 3**G**). These findings demonstrate that the high local GTF concentration created by clustering at the enhancer - and not the global nuclear GTF abundance -is the better determinant of transcription activity.

### Combined perturbations reveal synergy between promoter-enhancer interactions and activities of GTFs

Our discovery that GTF clusters form at distal enhancers, and that more frequent transcription bursting is associated with increased promoter proximity to such GTF clusters suggests a possible mechanism for how distal enhancers control transcription. Specifically, we hypothesize that enhancer-promoter communication might facilitate certain steps in the multi-step cascade that underlies burst initiation and that involves GTF activities. At the same time, some steps of this cascade might also occur independently of promoter-enhancer interactions, and might be accelerated by GTF activities alone.

Intuitively, if enhancer-promoter interactions and activity of a GTF accelerate different steps of the cascade (e.g. steps A and B), perturbing each process separately will add an additional delay in the initiation of bursts, *τ*_A_ and *τ*_B_, by slowing down steps A and B, respectively. If then both enhancer-promoter interactions and GTF activity are simultaneously perturbed, the total delay to complete the cascade will be slowed down by the sum of the individual delays, *τ*_A_+*τ*_B_. If steps A and B are rate-limiting, the frequency of transcription bursts will be decreased roughly proportionally to the delay time; the total reduction in transcription activity will then be the *additive* effect of the individual perturbations. Alternatively, if enhancer-promoter interactions and GTF activity both accelerate a single step of the cascade, each perturbation proportionally slows down that step. When both processes are simultaneously disrupted, the combined result is a *multiplicative* effect of the individual perturbations, with a total delay time ∼ *τ*_A_*τ*_B_/(*τ*_A_+*τ*_B_).

We further explore these ideas by numerical studies of the behavior of a two-step promoter (*32, 33*) (Supplementary Fig. 6**A**, Supplementary Note), regulated according to the following schemes: (1) the two steps are independently regulated by two different factors; (2) two factors simultaneously regulate one of the two steps. For a range of parameters, our results show that combined perturbations of the two regulatory factors result in (sub)-additive and (super)-multiplicative effects for schemes (1) and (2), respectively (Supplementary Fig. 6**B**). These two distinct predictions then provide a means for discerning whether enhancer-promoter interactions and a particular GTF activity might work together during steps of the cascade.

To further understand to what extend the burst initiation cascade might be controlled by promoter-enhancer interactions, we performed double perturbation experiments, simultaneously depleting Rad21 – to disrupt promoter-enhancer interactions – and perturbing selected GTFs. Double perturbations of Rad21 and XBP, Gtf2f1, Taf1, or Med19 all showed further reductions in the on-off bursting kinetics (Fig.4 **A-D**, Supplementary Fig. 7). Bursting frequencies were reduced ∼1.4-4.5-fold, while non-bursting fraction was increased ∼4-9-fold, for an overall decrease in the probability of observing a burst of 6-38-fold. At the same time, although bursts were very infrequently observed, when they occurred burst amplitude and duration were only modestly (up to 33% max, with most perturbations showing ≤15% changes) affected (with the exception of Rad21-Taf1 double perturbation, which shows a 2-fold increase in amplitude).

**Figure 4.**
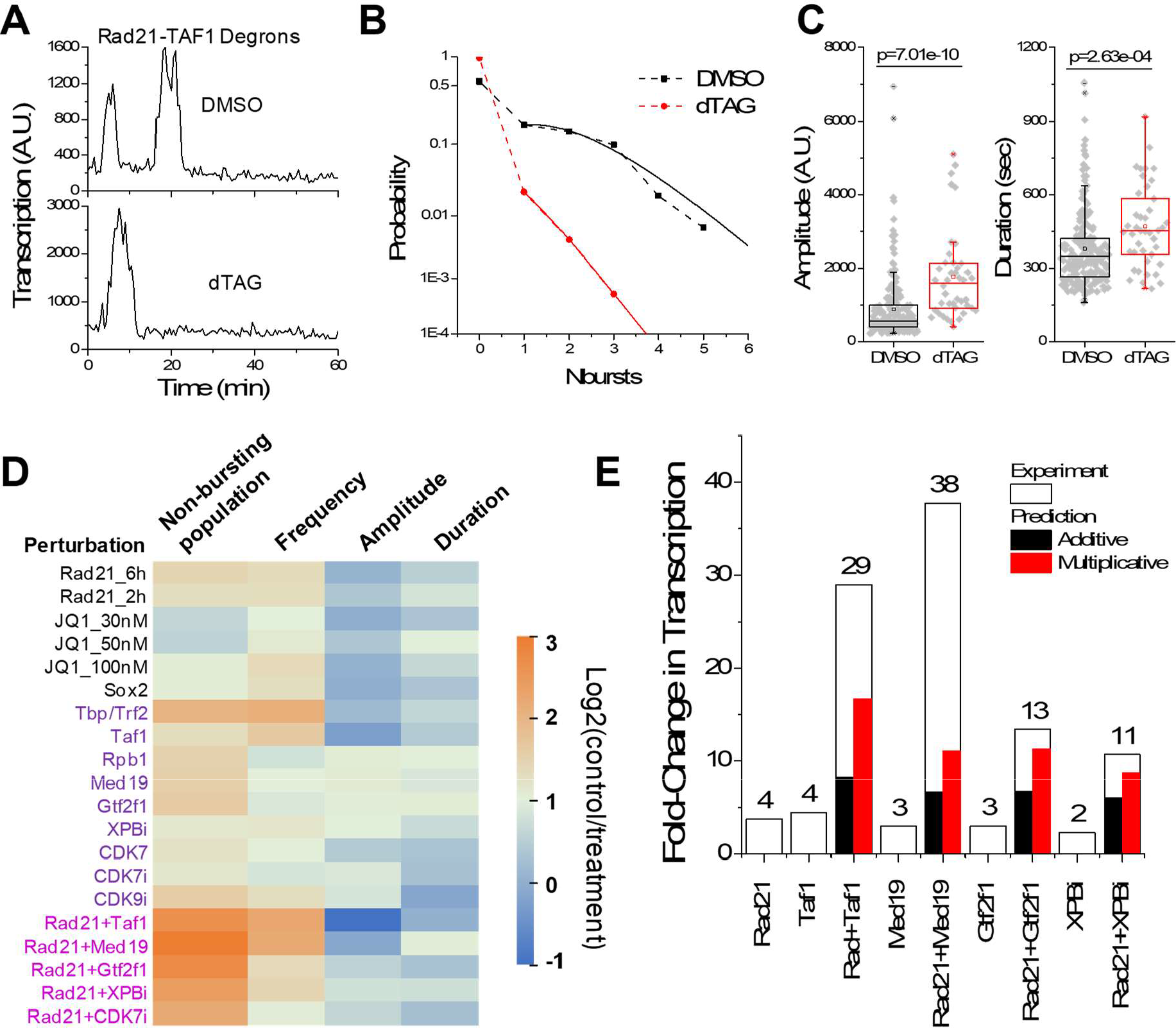
Synergies between promoter-enhancer interactions and activities of GTFs. (**A**) Two intensity traces of transcription sites from DMSO- and dTAG-treated Rad21-Taf1-FKBP^F36V^ KMG cells. (**B**) Probability of observing certain number of bursts in an 1 hr observation time, in DMSO-(*n*=135 traces from 152 cells) and dTAG-treated (*n*=51 traces from 935 cells) Rad21/Taf1-FKBP^F36V^ KMG cells. Solid lines: fits to Poisson distribution, for the data with *n*_bursts_≥1. The resulting two parameters (A, *r*) are: DMSO (0.56 and 1.69), dTAG (0.086 and 0.38). (**C**) Box plots of burst amplitude and duration. Each gray dot represents one burst. Data points are from two independent experiments with total n= 189 and 42 bursts for DMSO and dTAG, respectively. p-values are calculated based on a Wilcoxon rank-sum test. The data include two independent experiments. (**D**) Heat-map of fold-change of bursting parameters (non-bursting population, frequency, amplitude and duration) with double perturbations in KMG cells, including double dTAG degrons (Rad21 and Taf1, Med19, or Gtf2f1), dTAG combined with inhibitors (Rad21 and XPBi, or CDK7i). Data are normalized by Log2 (control/treatment). (**E**) Fold-change of control/treatment in single and double perturbations. Black and red boxes are the predicted additive and multiplicative fold-change, respectively. The white boxes show the experimental data, which are larger than both predicted additive and multiplicative effects. The data are from 2-3 independent experiments.

Moreover, quantification of the total reduction in transcription activity shows super-additive effects: the combined perturbations result in multiplicative or even super-multiplicative loss of activity (Fig. 4**E**, Supplementary Fig. 6**C**). These results indicate high overlap between the cohesin-mediated promoter-enhancer interactions and the activities of the selected GTFs, likely reflecting the action of both processes on the same step(s) of the multi-step burst initiation cascade.

## Conclusion

Our study provides a mechanistic explanation for the previously enigmatic phenomenon of enhancer control of transcriptional bursting. Our results reveal that the initiation of transcriptional bursts is a multi-step cascade, comprising several steps of the early phases of the Pol II transcription cycle. Bringing a promoter to within ∼100-200 nm of its distal enhancers accelerates steps in the burst initiation cascade. These close spatial encounters synergize with GTF activities to control the on-off bursting kinetics of the target promoter. For some GTFs, their activities might be physically facilitated by increasing the local GTF concentration in the vicinity of the promoter -as the promoter frequently interacts with clusters of GTFs that form at the enhancer. The dynamic range of productive promoter-enhancer communication thus can extend beyond the molecular-scale bridges posited by DNA looping models, to the mesoscopic size of these newly-described nano-scale nuclear environments.

## Acknowledgments

We thank Luke Lavis (HHMI/Janelia) for dye-labeling reagents, and Alessio Ciulli, Dario Alessi, and the MRC PPU Unit (University of Dundee) for HaloPROTAC-E compound.

## Author contributions

L.C. and J.L. generated mESC cell lines. L.C. performed live imaging experiments. L.C., C.D., and J.L. performed OligoPAINT FISH experiments. L.C., C.D., J.L., and A.P. analyzed data. A.P. wrote the manuscript with input from all the authors.

## Data and code availability

Data that support the findings in the paper and custom-written analysis code will be made available in an online depository upon publication of the peer-reviewed article.

**Supplementary Figure 1.**
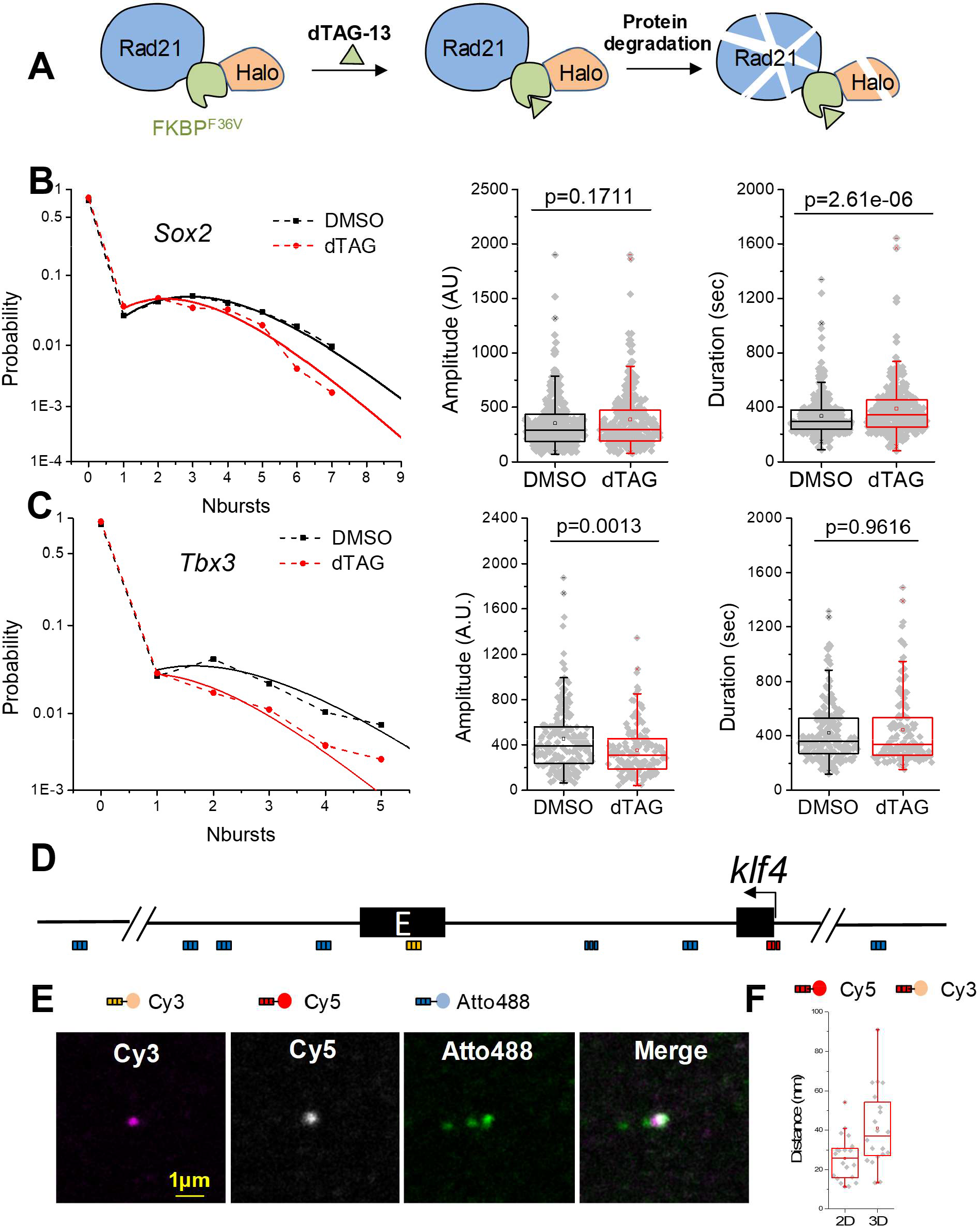
dTAG targeted degradation system in mESCs, Rad21 controls of *Sox2* and *Tbx3* on-off bursting, and high-resolution DNA OligoPAINT FISH. (**A**) Schematic of targeted protein degradation by dTAG-13. Endogenous Rad21 protein is fused with FKBP^F36V^ and Halo-tag. After dTAG-13 treatment, by staining with the Halo-tag, Rad21 protein can be measured (Figure 1**B**). (**B**) *Sox2* burst probability distribution, amplitude and duration in DMSO- and dTAG-treated Rad21-FKBP^F36V^ SMG cells. The data are from 2 independent experiments with total *n*=494 bursts from 211 traces in 465 cells, and 368 bursts from 209 traces in 572 cells, in DMSO and dTAG respectively. (**C**) *Tbx3* burst probability distribution, amplitude and duration in DMSO- and dTAG-treated Rad21-FKBP^F36V^ TMG cells. The data are from 2 independent experiments with total *n*= 188 bursts from 108 traces in 481 cells, and 131 bursts from 100 traces in 757 cells, in DMSO and dTAG respectively. (**D**) Schematic of the DNA OligoPAINT FISH probe sets, including plus_1kb (promoter), +55kb (enhancer), and additional regions (−60kb, +15kb, +30kb, +69kb, +76kb and +86kb) used as a reference to identify the *Klf4* locus. (**E**) Representative confocal images of DNA FISH signals. Enhancer probes are imaged by Cy3, promoter probes are imaged by Cy5, and the reference probes are imaged by Atto 488. (**F**) The 2D and 3D distances are measured between promoter(+1kb) simultaneously tagged by Cy5 and Cy3, indicating the precision of DNA FISH measurement.

**Supplementary Figure 2.**
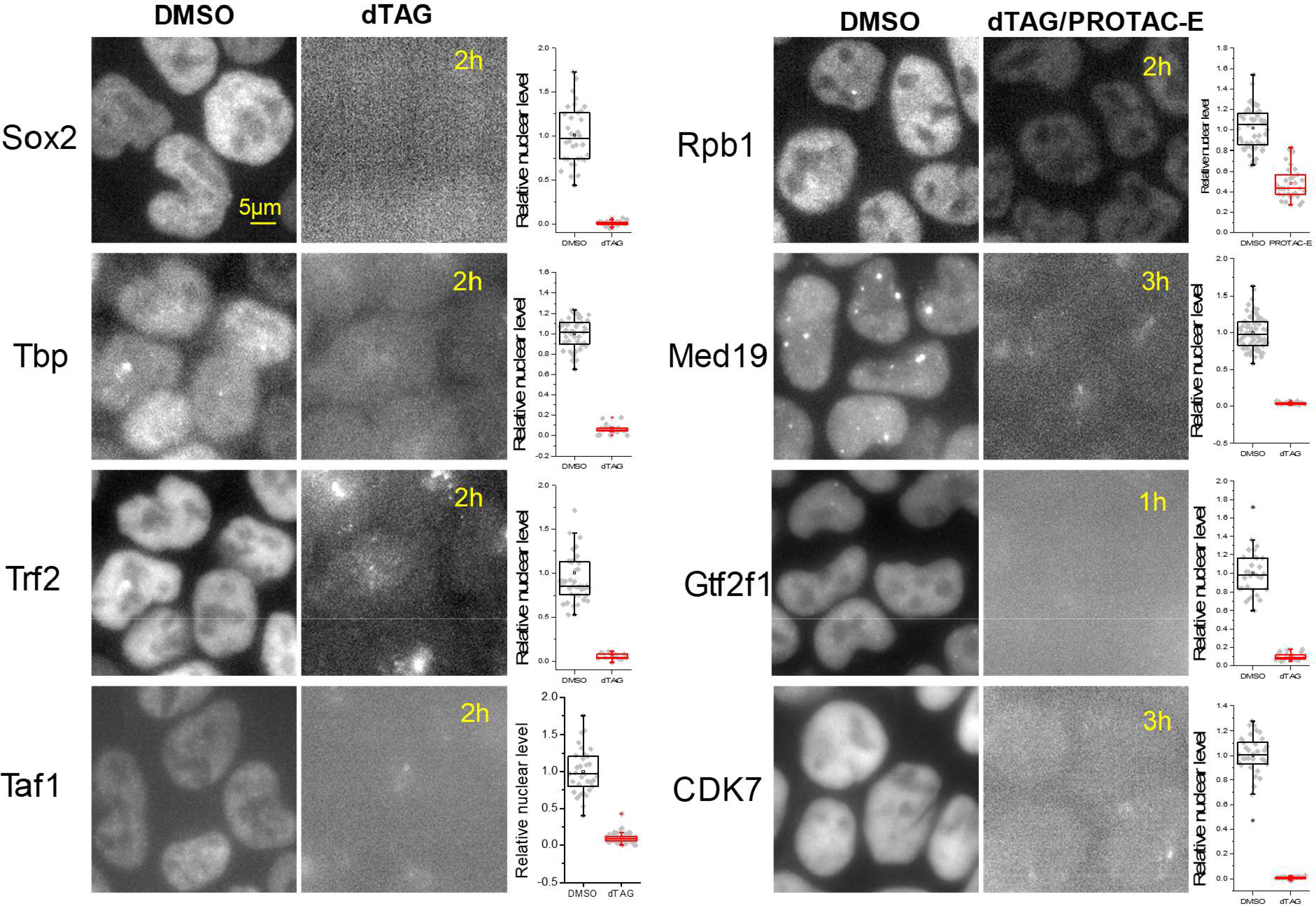
Targeted protein degradation of different factors participating in the transcription process. (Left panels) Representative Epifluorescence microscope images of nuclear Halo-Sox2, -Trf2, - Taf1, -Rpb1, -Med19, -Gtf2f1 and -CDK7, and SiR-Tbp in DMSO- and -dTAG or HaloPROTAC-E (Rpb1) treated cells. Treatment proceeded for the indicated number of hours (1-3 hr). (Right) Quantification of relative nuclear protein levels in the corresponding samples. Data are normalized to DMSO.

**Supplementary Figure 3.**
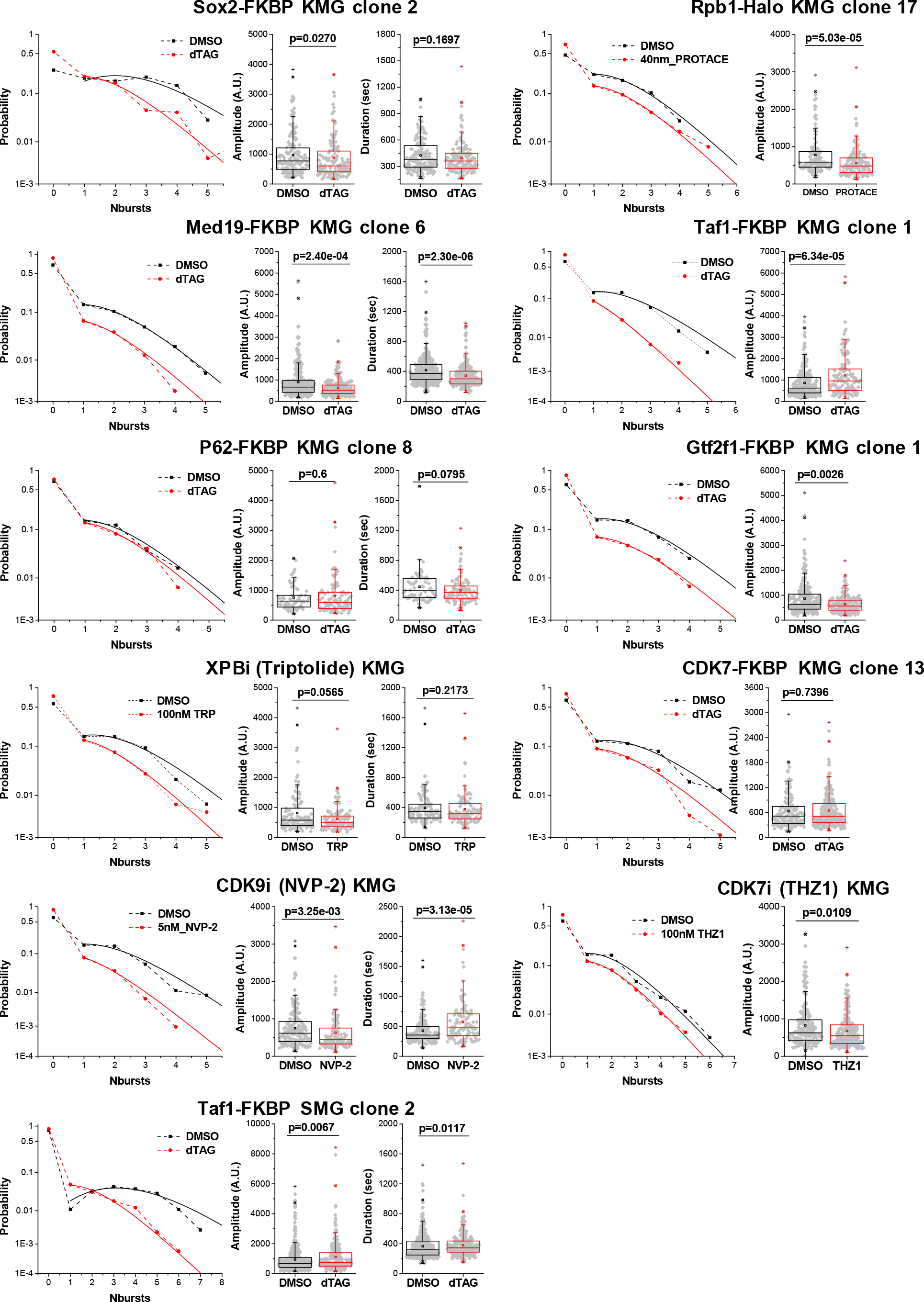
Transcriptional bursting kinetics is regulated by RFs (Sox2 and Med19), Rpb1 and GTFs (Gtf2f1, p62, CDK7 and XPB). Probability of number of bursts observed in 1 hr, as well as burst amplitude and duration, shown for different perturbations in KMG cells. Perturbations shown are dTAG targeted degradation (Rad21, Sox2, Tbp-Trf2, Taf1, Med19, Gtf2f1 and CDK7), Halo-PROTAC-E induced partial degradation (40 nM) of Rpb1, and non-saturating concentrations of inhibitors [100 nM THZ1 (CDK7i) and triptolide (XPBi), 5 nM NVP-2 (CDK9i)]. Similar results are also obtained with a Taf1 degron in SMG cells. The obtained fitting parameters A and *r*, as well as the total number of traces and cells analyzed are listed in Supplementary Table 2.

**Supplementary Figure 4.**
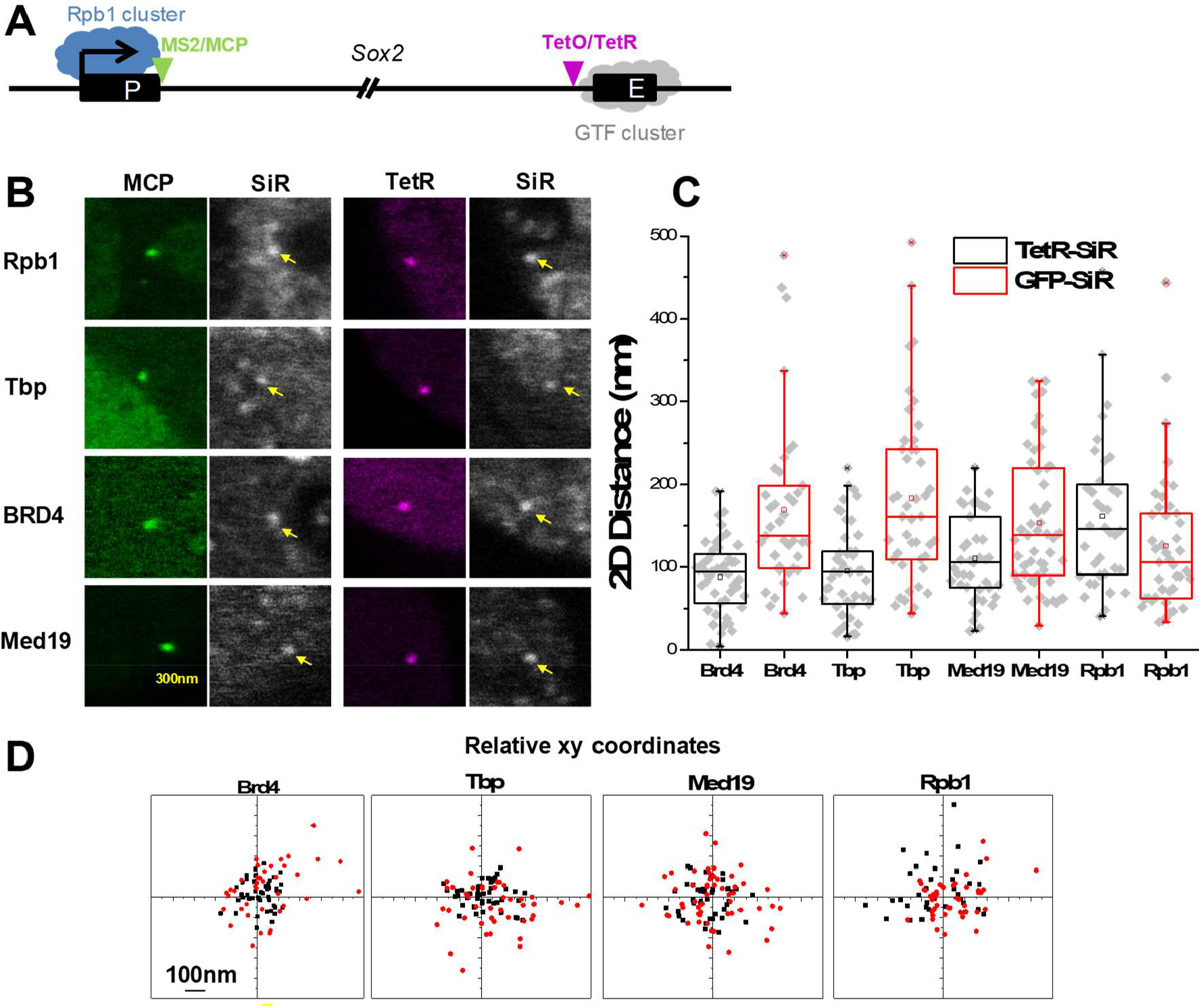
TBP, Brd4 and Med19 clusters form at the *Sox2* +110 kb downstream enhancer. (**A**) Schematic of edited *Sox2* locus. The nascent *Sox2* RNA is labelled by 24×MS2/MCP at the 3’ UTR. 48×TetO is integrated near the enhancer region (SCR), which can be visualized by TetR-Halo. (**B**) Representative live-cell confocal images show co-localized: MCP-mNeonGreen(green) and SiR-Rpb1, -Tbp, -Med19 and -Brd4 foci (left) and TetR-Halo (magenta) and SiR-Rpb1, -Tbp, -Med19 and -Brd4 foci (right). The yellow arrows point to the clusters. (**C**) Relative TetR- and MCP-Brd4, -Tbp, -Med19 and -Rpb1 2D distances, (mean ± SD): Brd4 88 ± 43(169 ± 102)nm, Tbp 95± 54(183 ± 105)nm, Med19 110± 61(153 ± 81)nm, Rpb1 161 ± 91(125 ± 83)nm, n = 47(39), 46(44), 41(55), 36(43), TetR-SiR (MCP-SiR) respectively. Each point corresponds to the distance of a single transcription site. All data points are from two independent experiments. (**D**) Relative *xy* coordinates corresponding to the points in (**C**).

**Supplementary Figure 5.**
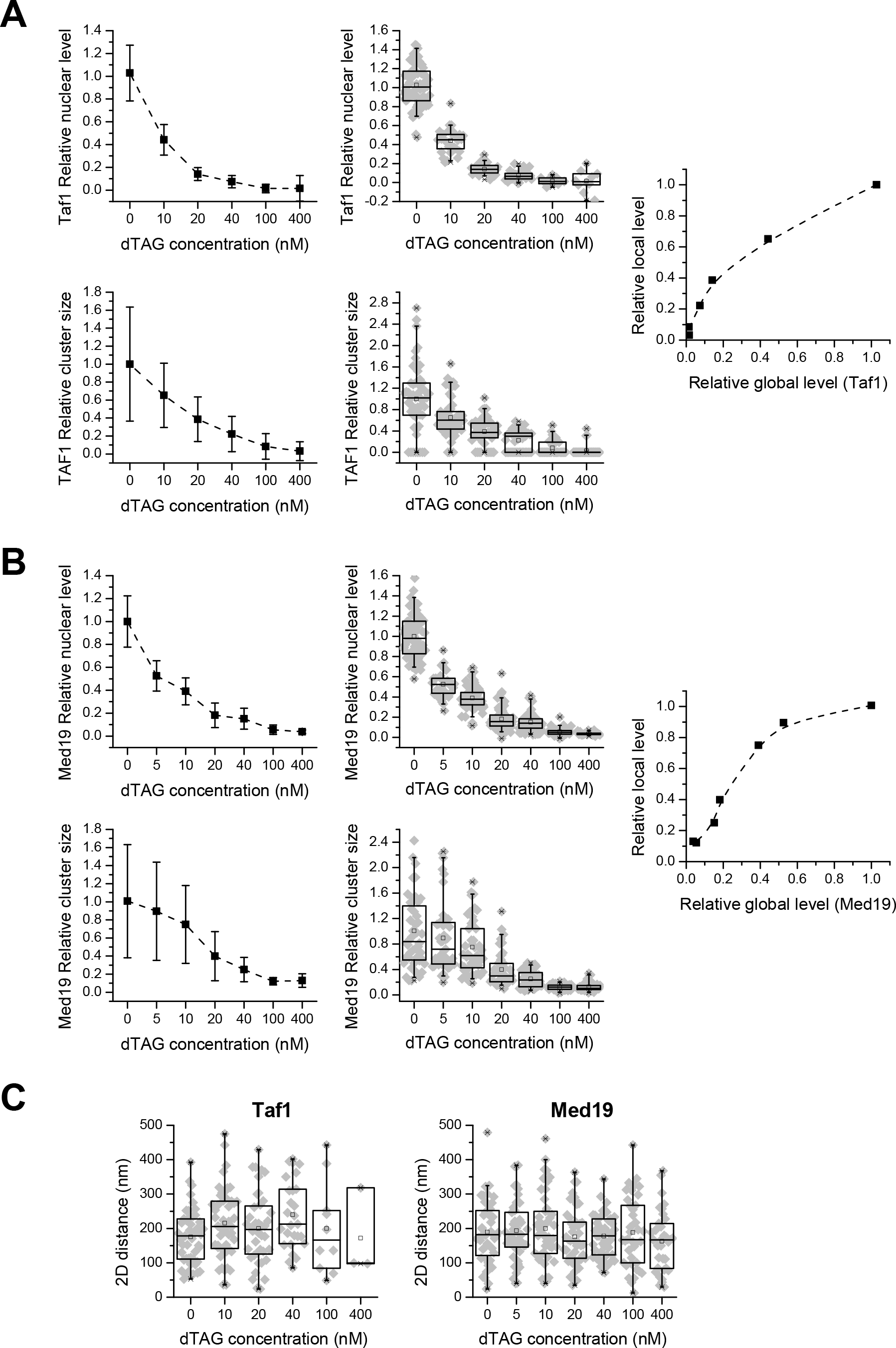
Modulation of global and local Taf1 and Med19 levels. (**A**) Upper: Realtive Taf1 nuclear level normalized to DMSO (dTAG=0) upon dTAG titration. Line plot shows with mean ± SD: 1 ± 0.24, 0.44 ± 0.13, 0.14 ± 0.06, 0.07 ± 0.05, 0.015 ± 0.04 and 0.015 ± 0.11, in dTAG = 0, 10, 20, 40, 100 and 400nM respectively. Each point in the box plot corresponds the relative level of a single cell, n= 71, 29, 31, 26, 11 and 10 respectively. Lower: Realtive Taf1 cluster size normalized to DMSO upon dTAG titration. Line plot shows with mean ± SD: 1 ± 0.63, 0.65 ± 0.36, 0.38 ± 0.25, 0.22 ± 0.19, 0.08 ± 0.14 and 0.03 ± 0.1, in dTAG = 0, 10, 20, 40, 100 and 400nM respectively. Each point in the box plots corresponds the relative cluster zise of a single transcription site, n= 55, 51, 49, 45, 33 and 34 respectively. The right line plot shows relative nuclear-cluster level (global-local) relationship based on the left data. (**B**) Upper: Realtive Med19 nuclear level normalized to DMSO upon dTAG titration. Line plot shows with mean ± SD: 1 ± 0.22, 0.53 ± 0.13, 0.39 ± 0.12, 0.18 ± 0.11, 0.15 ± 0.09, 0.06 ± 0.04 and 0.04 ± 0.16, in dTAG = 0, 5, 10, 20, 40, 100 and 400nM respectively. Each point in the box plot corresponds the relative level of a single cell, n= 66, 30, 70, 71, 65, 31 and 20 respectively. Lower: Realtive Med19 cluster size normalized to DMSO upon dTAG titration. Line plot shows with mean ± SD: 1 ± 0.63, 0.89 ± 0.54, 0.75 ± 0.43, 0.4 ± 0.27, 0.25 ± 0.13, 0.12 ± 0.04 and 0.13 ± 0.07, in dTAG = 0, 5, 10, 20, 40, 100 and 400nM respectively. Each point in the box plot corresponds the relative cluster size of a single transcription site, n= 48, 45, 45, 45, 45, 38 and 35 respectively. The right line plot shows relative nuclear-cluster relationship based on the left data. (**C**) Realtive Klf4 transcription site and cluster 2D distances upon dTAG titration corresponding to (**A** and **B**). Taf1 (mean ± SD): 175 ± 79nm, 215 ± 100nm, 199 ± 104nm, 239 ± 98nm, 199 ± 133nm and 171 ± 126nm, n= 47, 48, 40, 27, 10 and 3 respectively. Med19 (mean ± SD): 189 ± 90nm, 193 ± 88nm, 199 ± 102nm, 175 ± 83nm, 178 ± 62nm, 188 ± 102 and 163 ± 87nm, n= 48, 45, 45, 45, 45, 38 and 35 respectively. Each point corresponds to the distance of a single transcription site. All data points are from two independent experiments.

**Supplementary Figure 6.**
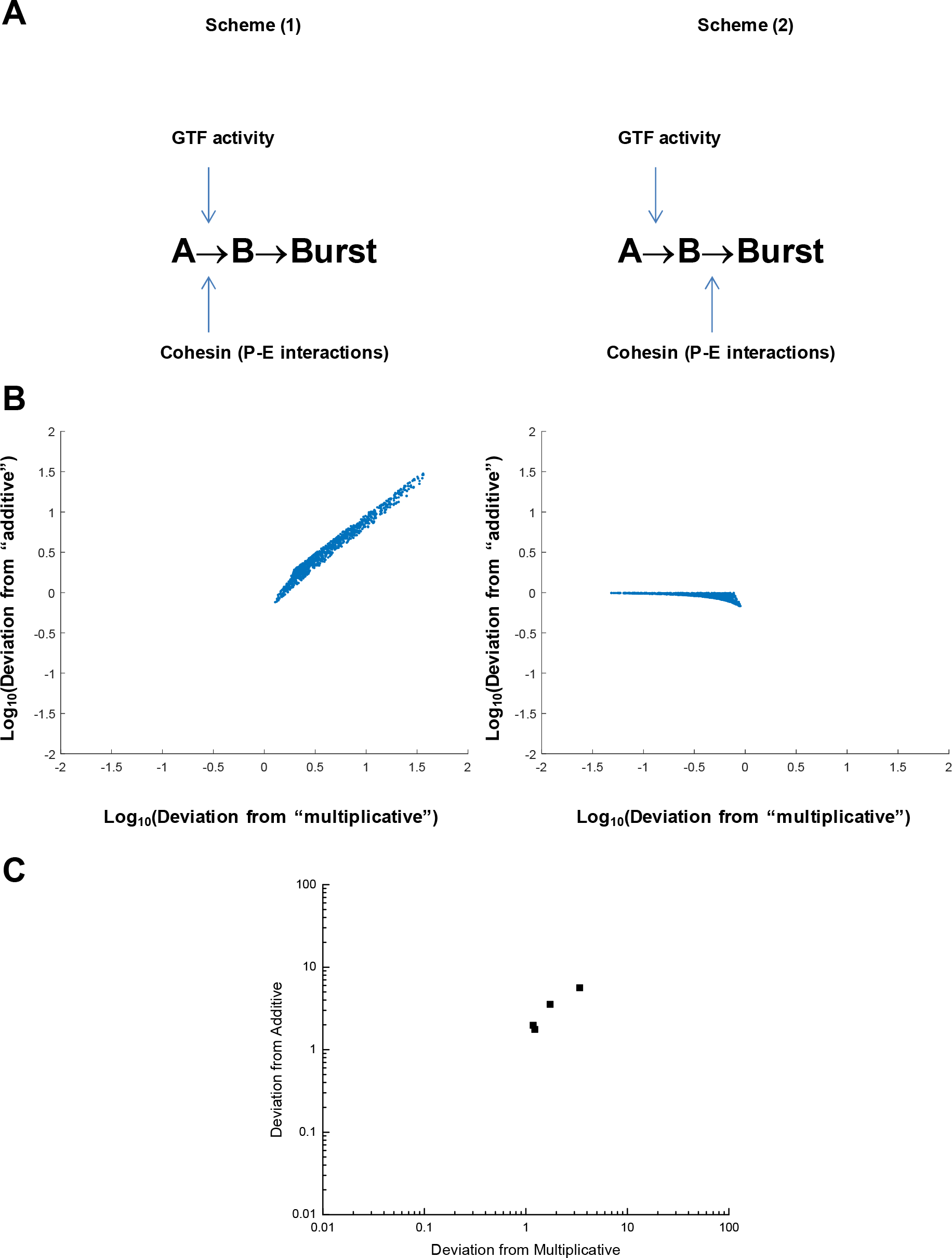
Numerical studies of the behavior of a two-step promoter. (**A**) Two different regulatory schemes are considered. In Scheme (1), a GTF activity and cohesin-mediated enhancer-promoter interactions act together to accelerate the first step of the promoter cascade. In Scheme (2), the GTF and cohesin activities independently accelerate the two different steps of the cascade. (**B**) Graphs show the deviation from additive and multiplicative behavior, observed when the GTF and cohesin activities are disrupted, individually and in combination. See Supplementary Note for details. (**C**) Deviations from additive and multiplicative behaviors for the experimental data in Fig. 4**E**.

**Supplementary Figure 7.**
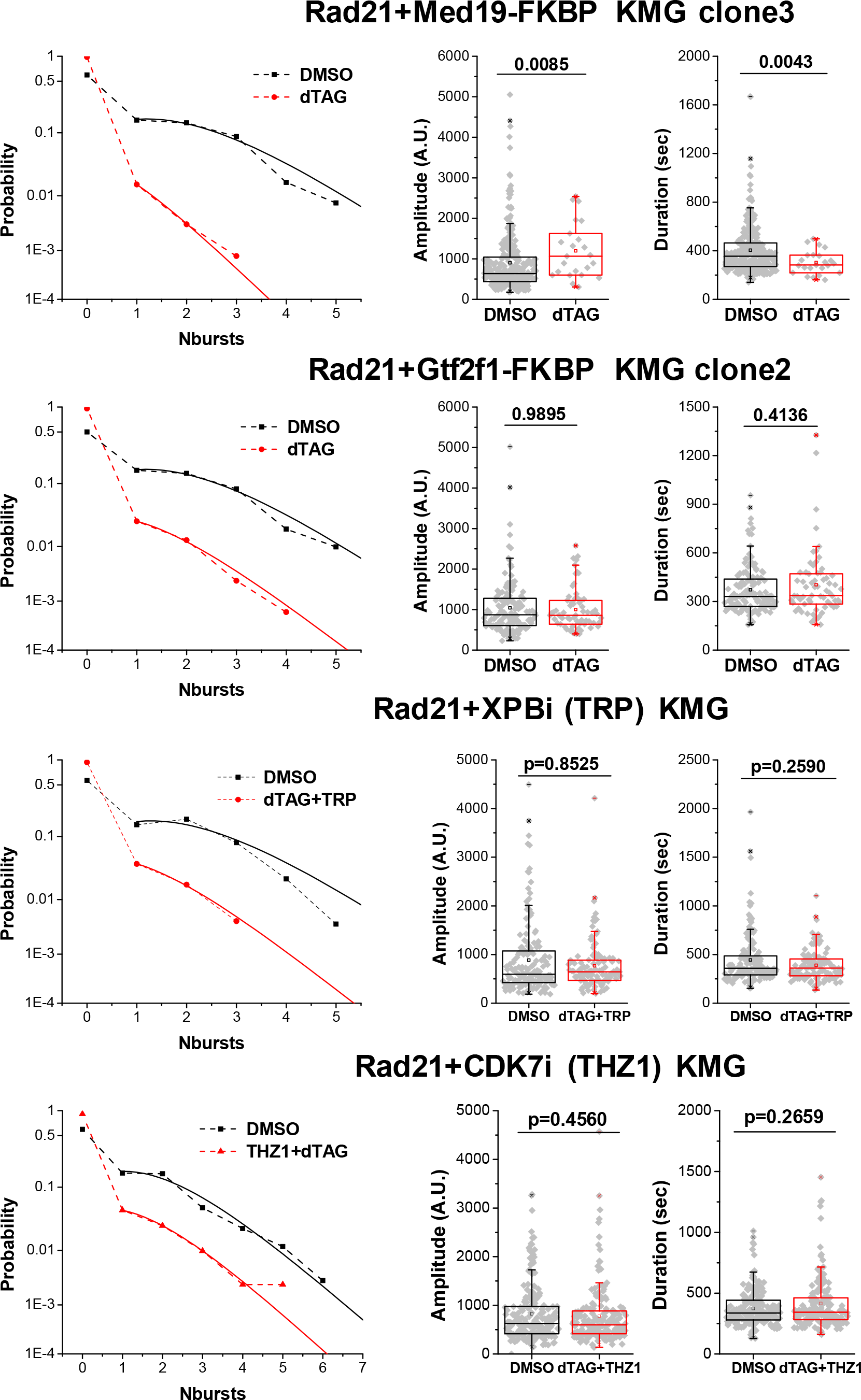
Combined perturbations of Rad21 and GTFs. Probability of number of bursts observed in 1 hr, as well as burst amplitude and duration, shown for double perturbations in KMG cells. Perturbations shown are double dTAG targeted degradation (Rad21 and Med19, or Gtf2f1), and dTAG targeted degradation of Rad21 combined with small-molecule inhibitors (Rad21 and XPBi, or CDK7i). The data are from two independent experiments. The obtained fitting parameters A and *r*, as well as the total number of traces and cells analyzed are listed in Supplementary Table 2.

## Materials and Methods

### Cell lines

Mouse embryonic stem cell lines were derived from Bruce 4 mESCs (Millipore Sigma CMTI-2; murine strain C57/BL6J, male – species/sex verified by karyotyping, no additional cell line authentification performed). KMG mESCs contain 24 × MS2 cassettes integrated in the 3′-UTR of both *Klf4* alleles and stably express MCP-mNeonGreen. SMG mESCs(*11*) contain 24 × MS2 cassettes integrated in the 3′-UTR of one *Sox2* alleles and stably express MCP-mNeonGreen.

### Cell culture

All mESC lines were maintained in 2i medium with appropriate selection drugs, containing DMEM (Thermo Fisher Scientific 10313021), 15% fetal bovine serum (FBS, R&D Systems, S10250), 0.1 mM 2-mercaptoethanol (Thermo Fisher Scientific 21985023), 1× GlutaMAX™ Supplement (Thermo Fisher Scientific 35050079), 1× MEM nonessential amino acids (Thermo Fisher Scientific 11140076), 1000 U/mL LIF (Millipore ESG1107), 3 µM CHIR99021 (Millipore 361559) and 1 µM PD0325901 (Axon Medchem 1408) and 100U/mL Penicillin-Streptomycin (Thermo Fisher Scientific, 15140122) on a 0.1% gelatin-coated dish at 37°C and 5% CO2, in a humidified incubator.

### CRISPR/Cas9 gRNA cloning and validation

gRNAs were designed using an online tool (CRISPR gRNA Design tool – ATUM, https:www.atum.bio/). For gRNA cloning, oligo pairs were annealed and ligated into BbsI -digested espCas9 plasmid (Addgene 71814). For validating the efficiency of gRNAs, 250ng of espCas9-gRNAs were transfected into 1×10^4^ mESC cells using Lipofectamine 2000 (Invitrogen 11668019). Genomic DNA was extracted 3 days post-transfection using QuickExtract DNA Extraction Solution 1.0 (Lucigen Corporation QE09050). To test the cutting efficiency, surveyor assays were performed. Briefly, DNA samples were PCR-amplified by KOD One™ PCR master Mix (TOYOBO KMM-101) using site-specific primers, the PCR products were denatured by heating-up and then cooled down to form heteroduplexes. Mismatched duplexes were then cleaved by T7 Endonuclease I (NEB M0302S) and cleavage products were detected by gel electrophoresis. All gRNAs used for generating cell lines are listed in Table 1.

### Generation of mESCs with 24 × MS2 integration at the *Klf4* and *Tbx3* 3′ UTR and stable expression of MCP-mNeonGreen. Targeting vector construction

The 24**×** MS2 was integrated before the stop codons of the mouse *Klf4* and *Tbx3*. The targeting vectors were assembled in two steps. First, vectors were synthesized containing part of the *Klf4* or *Tbx3* coding sequence (878 bp or 1 kb) as a HA-L, a part of T2A-hygromycin and followed by part of the 3’UTR sequence (990 bp or 1 kb) as a HA-R, resulting in pUC57-HA-L-T2A-HygR-HA-R. Next, a 24**×**MS2 cassette was cut from pCR4-24**×**MS2SL-stable (Addgene 31865) and pasted into pUC57-HA-L-T2A-HygR -HA-R, resulting in the donor vectors pUC57-HA-L-T2A-HygR -24**×**MS2-HA-R.

Bruce 4 mESCs (1×10^6^) were transfected with 10µg donor vector and 0.6µg espCas9-Klf4 using Lipofectamine 2000 (Invitrogen 11668019). After 3 days, cells were subjected to Hygromycin B selection. Individual colonies were picked and expanded. Clones were screened by genotyping and single clones with bi-allelic integrations were selected for stable integration of MCP-mNeonGreen. Cells were transfected with 10 µg pPB-LR5-CAG-MCP-mNeonGreen-IRES-Neo and 1 µg pCMV-hyPBase vectors using Lipofectamine 2000 (Invitrogen 11668019). After incubation for 2 days, cells were subjected to 400 µg/ml G418 (Sigma G8168) selection.

Individual colonies were picked, expanded, and imaged. A single clone (Klf4 MCP clone 7, dubbed “ KMG”) was selected for all further experiments. Similarly, a single clone (*Tbx3* MCP clone 7, dubbed “ TMG”) was selected for all further experiments.

### Generation of genome-edited mESCs for visualization and targeted degradation of endogenous protein factors

The donor plasmids used to target each endogenous mouse protein factor were synthesized in GenScript. Two homology arms (750bp left arm and 750bp right arm) of the targeted genes and the sequence encoding FKBP^F36V^–Halo or FKBP^F36V^–SNAP was synthesized and cloned into the pUC57mini vector. PAM sequences of the desired gRNAs were mutated to avoid re-editing. Rad21, TRF2, GTF2F1, p62, and CDK7 were tagged at their C-termini, while TBP, TAF1, GTF2E2 and Med19 were tagged at their N-termini.

To generate knock-in cell lines, KMG mESCs (1×10^6^) were transfected with 10 μg donor vector and 0.6 μg of the corresponding espCas9-gRNA vector. 7 days after transfection, the cells were labeled with 0.3 μM Halo-or SNAP-dye and followed by immediate fluorescence-activated cell sorting (FACS) to collect Halo-positive or SNAP-positive cells. Once we obtained nearly 100% positive pools, the cells were diluted, and single clones were picked and expanded. Insertion of tags was confirmed by PCR. The primers for genotyping are listed in Table 1. Homozygous clones showing nuclear staining and bright transcription sites were selected for all further experiments.

### Generation of KMG mESCs with genomic deletions

To delete the sequences between the *Klf4* promoter and its distal downstream enhancer, several gRNAs were designed in the downstream region of *Klf4* and cloned into pspCas9-2A-puro (PX459, Addgene 62988). Before the day of transfection, 2×10^5^ cells were seeded on a 0.1% gelatine-coated 6-well plate. Cells were transfected with 1µg of two Cas9-gRNAs using Lipofectamine 2000 (Invitrogen 11668019). Next day, the cells were transferred to a 10cm dish and subjected to selection with 1 µg/ml Puromycin (Thermo Fisher Scientific A1113803) for 3 days. After 7 days, individual clones were picked and transferred to a 96-well plate. Individual clones were genotyped by PCR and DNA sequencing. Homozygously targeted clones were selected for further experiments. The gRNAs and primers for genotyping are listed in Table 1. Klf4-1K-g5 and Klf4-42K_g5 were used to generate 42K deletion cells.

### Preparation of mESCs for live-cell imaging and targeted degradation of selected protein factors

Cells were seeded onto 8-chamber coverglass (Nunc™ Lab-Tek™ II Chambered Coverglass, 155409) coated with 5 µg/ml laminin (BioLamina LN511), in 2i media and appropriate drugs. For all experiments, cells were grown overnight in the chambers. Protein degradation was induced by treating cells with a final concentration of 400 µM dTAG-13 (Sigma SML2601) for 2 hrs or 6 hrs (2 hrs: Sox2, Taf1, Tbp/Trf2, Gtf2f1, CDK7; 6 hrs: Rad21 and Med19). 0.04% v/v DMSO was used as control. Before imaging, cells were labeled with the relevant Halo-dye (Promega, Janelia Fluor® HaloTag® Ligands) or SNAP-dye (SNAP-Cell® 647-SiR, NEB S9102S) with media containing 0.3 µM dye for 10 min, at 37°C, followed by three times rinsing with new media. Rpb1 protein degradation was induced by treating cells with 40 nM Halo-PROTAC-E(*34*) for 2 hrs and 5 nM Halo-dye for 1 hr. For THZ1 and Triptolide treatment, cells were treated by 100 nM for 1hr. For combined dTAG and inhibitor treatment, cells were treated by dTAG-13 for 6 hrs and 100 nM inhibitor for 1hr.

### Design of OligoPAINT DNA FISH probes

The oligonucleotides of the primary hybridization probes (referred to hereafter as primary probes) were designed by concatenating the following sequences from 5’ to 3’: 1) a unique 20-nt readout sequence for each targeted genomic region, complementary to the last 20 nt of the readout probes; 2) a 40-nt target sequence that allows hybridization to targets spanning around 3kb of the genomic region of interest. The readout sequences were designed according to previous studies (*35*). Readout probe is a 51-nt oligonucleotide containing on one end complementary sequence to primary probes and on the other end complementary sequence to the imaging probe. The imaging probes contain 21-nt oligonucleotide sequence complementary to the readout probe, and are modified with Atto 488, Cy3 or Cy5 dye on the 5’ end.

The 40-nt target sequences complementary to the genomic region of interest were selected according previous designs (*29, 35*). For a genome region of interest, we masked repeats to avoid off-targeting and get all potential 40-nt sequences (starting at each possible base in the targeted region). Then the probes were picked based on the following criteria: melting temperature in the range 65 – 80 °C and GC content 30 – 80%. The probes selected have no overlap with adjacent probes. We designed target sequences of *Klf4* promoter (plus 1kb) and enhancers (downstream 55kb and 69kb) with the following coordinates: chr4:55527143-55533466 (mm10), chr4:55488180-55491680(mm9) and chr4:55476500-55479600 (mm9). We also designed probes for 6 regions spanning from 60kb upstream to 110kb downstream of *Klf4*, which were used as reference to identify the gene locus. Each set of primary probes targeted around 3kb and contained ∼50 probes, which was assigned a unique readout sequence as described above.

The oligo pools of primary probes were synthesized by Integrated DNA Technologies (IDT).

### Preparation of cells for OligoPAINT DNA FISH experiments

Cells were seeded onto 5 µg/ml laminin (BioLamina LN511) coated 8-chamber coverglass (Nunc™ Lab-Tek™ II Chambered Coverglass, 155409) in 2i media and appropriate drugs. Next day, the cells were treated with dTAG-13 to induce targeted protein degradation or with small-molecule chemical inhibitors. For FISH experiments(*36*), the samples were fixed at room temperature (RT) in 4% formaldehyde (Thermo Scientific, 28906) in PBS for 10 minutes. Our fixation buffer (4% FA) is completely methanol free to avoid the nuclear shrinkage at the time of fixation. After three washes, the fixed cells were treated with 0.1 % (w/V) NaBH4 (Sigma Aldrich) for the reduction of excess formaldehyde for 7 minutes at RT followed by three PBS washes. Then we performed a two-step permeabilization as follows:(1) treated with 0.5% (v/v) Triton-X (Sigma Aldrich, T8787) in PBS for 10 minutes at RT and (2) treated with 0.1 M Hydrochloric acid (Sigma Adrich) for 5 minutes at RT. After three rounds of PBS washes following each step, we treated the cells with 0.1 mg/mL RNase A (Thermo Fisher Scientific, AM2271) in PBS for 45 minutes in 37 °C to avoid the cross hybridization of our primary probes with endogenous cellular RNA. Subsequently, the samples were incubated in prehybridization buffer (2× saline sodium citrate buffer (SSC) (Thermo Fisher Scientific, AM9763) and 50% formamide (Merck, S4117) for 30 minutes in RT. Next, cells in each well were incubated with 100 µL of hybridization buffer (2× SSC, 50% formamide and 10% w/v dextran sulfate (Sigma, D8906-50G) mixed with ∼1 µM total primary probes) for another 30 minutes at RT. Then the chamber was incubated at 86 °C on a metal block in a hybridization oven for 7.5 minutes.

Following this, the chamber was quickly transferred to 41 °C humidified incubator for 18-20 hours. In the meantime, the readout probes were annealed with corresponding imaging probes (Cy5/Cy3/Atto 488) in annealing buffer (1mM Tris-HCl pH 8.0, 0.1 mM EDTA, 5mM NaCl). Then, the samples were washed with washing buffer (2× SSC, 40% formamide) for 6 times and kept in RT for 30min. For second hybridization, each well was incubated with 100 µL hybridization buffer mixed with the annealed probes for 30min at RT. Thereafter, the sample was washed several times with washing buffer to remove the excess unattached imaging probe form the sample. Finally, samples were stored at 4 °C in 2× SSC until imaging.

### Live-cell imaging of nascent 24 × MS2 transcription dynamics

A wide-field epifluorescence microscope (Zeiss ZEN) was used to image nascent transcription dynamics. We used a 63× 1.4 NA oil-immersion objective lens, FITC filter and 4% intensity of an XCite EXACT fluorescence excitation source. We imaged an *xyz* volume ≈211×211×2.5-5 µm^3^ containing multiple cells, using 270nm *z* steps. Camera exposure times were 500 ms at each *z* position, resulting in 15-20 sec/volume total time. The samples were imaged for 1 hr at 30 s intervals. Live-cell imaging was performed at 37°C and 5% CO_2_ atmosphere.

### Live-cell imaging of RF clustering at single transcription sites

RF clusters were imaged with a Leica TCS SP8 confocal setup, featuring far-red-sensitive Avalanche-Photo-Diode detectors (Excelitas) and a white-light super-continuum laser excitation source (NKT Photonics SuperK EXTREME EXW-12). Imaging was performed with a 63× 1.20 NA water immersion objective lens (Leica 15506346 HC PL APO 63×/1.20 W CORR CS2) at 490nm and 648nm excitation. We scanned an *xyz* volume ≈3.84×3.84×3 µm^3^ around the transcription site, using 30.27 nm *xy* pixel size and 300 nm *z* steps. At each 300 nm *z*-slice, an *xy* scan was completed in ≈1.8 seconds, resulting in ≈20 sec/volume total time. Live-cell imaging was performed with the whole microscope enclosed inside a temperature-controlled box. An additional stage incubator (Tokai-Hit) was used to regulate temperature and atmosphere of the sample at 37°C and 5% CO_2_, respectively.

### Fixed-cell imaging of tagged genomic loci using OligoPAINT FISH

DNA oligoPAINT FISH imaging was performed at 24 °C in a Leica TCS SP8 confocal microscope equipped with 100× 1.44 NA oil immersion objective (Leica, HC PLAN APO), two SPCM AQR (Perkin Elmer) APD detectors (for Cy5 and Cy3 emission detection) and a HyD Reflected Light Detector (RLD) in gated mode (for Atto 488 emission detection). Oxygen scavenger solution (75mM HEPES-KOH pH7.5, 55mM K-glutamate, 0.9% w/v Glucose, 1mM Ascorbic Acid, 1mM Methyl-viologen, 1×gloxy) was used to protect signals from bleaching(*37, 38*). In exposure 1, OligoPAI4NT oligos with Cy5 and Cy3 imaging probes were excited with 551 nm and 649 nm laser light, respectively. The emissions were collected through a dichroic mirror (Shemrock 625 nm edge beamsplitter) in two APD detectors. To avoid further crosstalk between Cy3 and Cy5 emission, we used two band pass filters (Chroma ET595/50m for Cy3, Chroma ET690/50m for Cy5) in front of the two APDs, respectively. In exposure 2, OligoPAINT oligos with Atto 488 imaging probes were excited with 488 nm laser light and the emission was collected in a different path (band pass filter Chroma ET525/50m) on the HyD SMD detector. We used 10%, 1% and 5-7% of maximum laser power to excite Cy3, Cy5 and Atto 488, respectively, selected to avoid bleaching. We scanned an *xyz* volume ≈5.81×5.81×3 µm^3^, using 46 nm *xy* pixel size and 100nm *z* steps.

### Image Analysis

Image processing was performed using custom MATLAB 2010b and R2020a (MathWorks) routines, as well as Fiji (ImageJ 1.52a).

### Quantification of transcriptional bursting

Maximum intensity projection images were calculated for each *z* stack, of MCP-mNeonGreen images, and used to track the intensities of transcription sites during bursting. A particle tracking routine (*39*) was used for identifying transcription sites in each frame. The intensity trace of MCP-mNeonGreen was fitted to a trapezoid function (*11*), to obtain burst amplitude and duration. The distribution of number of bursts per total acquisition time was fitted to a Poisson distribution, with an additional term denoting the non-bursting population.

### 3D localization and distance measurements using OligoPAINT FISH

3D distances were measured using in-house MATLAB scripts. To account for chromatic aberrations resulting in apparent shifts between Cy3 and Cy5 images, we used 0.1 μm TetraSpeck bead (Invitrogen, 1000× diluted in PBS) to calculate the 3D offsets between the two colors. Images analyzed showed multiple Atto 488 spots, tagging 6 positions over the extended *Klf4* locus. The *xy* coordinates of the Cy3 and Cy5 spots were obtained by least-squares fitting to 2D Gaussian functions. The z coordinate was then obtained by fitting the intensity *z* profile centered on the *xy* coordinates to a 1D Gaussian function plus a linearly-varying background. The 3D distance between Cy3 and Cy5 was then obtained from relative the *xyz* coordinates. By labelling the same genomic region with both Cy3 and Cy5 probes (“ zero” distance control), we estimate distance precisions of 26 nm in *xy* and 36 nm in *z*., for a total 3D distance precision of ≈41 nm.

### Statistics

Experiments were replicated multiple times, as indicated in the respective figure legends. Box-plots with descriptive statistics were generated in Origin. Boxes indicate IQR (25–75% intervals) and the median line, whiskers indicate 1.5 × the IQR; ‘×’ symbols indicate 1% and 99% percentiles; and square symbols indicate the mean. For significance testing, Wilcoxon rank sum statistical tests were performed in MATLAB.

