## Supplementary Note for "Mechanisms of transcription control by distal enhancers from high-resolution single-gene imaging"

### NUMERICAL STUDIES OF TWO-STATE PROMOTER MODELS

#### 1. Definition of models.

We consider a two-state promoter model (*1, 2*), where the promoter transitions between two OFF states, $P_1$ and $P_2$, before it produces a burst. We further consider two regulatory schemes, (1) and (2) (Supplementary Note Fig. 1). In scheme (1), two factors, A and B, simultaneously accelerate the first step in the promoter cascade. In scheme (2), factors A and B accelerate the first and second step in the promoter cascade, respectively.

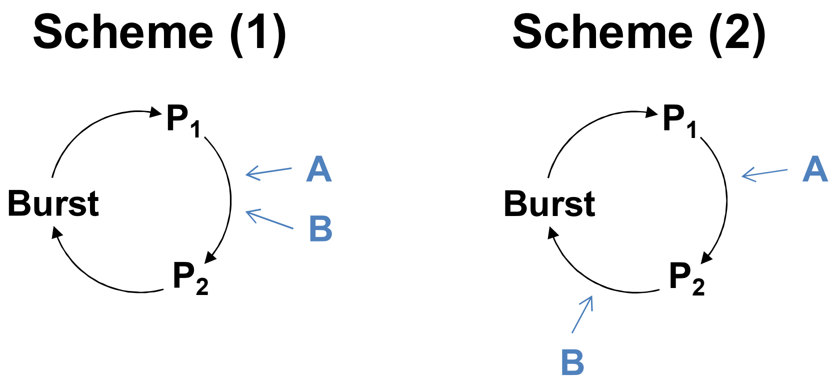


**Supplementary Note Figure 1. Two-state promoter models.**

Without factors A and B, the transition rates between states are $k_1$ and $k_2$ for $P_1 \rightarrow P_2$ and $P_2 \rightarrow$Burst, respectively. When factors A and B are present, these rates are accelerated accordingly (*2*):

$k_1' = k_1$ (a+b+1), $k_2' = k_2$ Scheme (1).

$k_1' = k_1$ (a+1), $k_2' = k_2$ (b+1) Scheme (2).

The total transcription bursting rate is given by: $1/k_{trxn}(a,b) = 1/k_1' + 1/k_2'$. Perturbations of factors A and B result in corresponding relative losses in transcription activity:

$R_A = k_{trxn}(a,b) / k_{trxn}(0,b)$ loss of A.

$R_B = k_{trxn}(a,b) / k_{trxn}(a,0)$ loss of B.

$R_{AB} = k_{trxn}(a,b) / k_{trxn}(0,0)$ loss of both A and B.

We define an 'additive" effect of loss of both factors A and B if $R_{AB} = R_A + R_B$, and a "multiplicative" effect if $R_{AB} = R_A R_B$. Similarly, we define parameters $\Delta_a = R_{AB} / (R_A + R_B)$ and $\Delta_m = R_{AB} / (R_A R_B)$, which quantify the deviation from "additive" and "multiplicative" behaviours, respectively.

For scheme (1), we obtain the following expressions for $\Delta_a$ and $\Delta_m$:

$$\Delta_a = \frac{\left(\frac{1}{k_1} + \frac{1}{k_2}\right)}{\left(\frac{1}{k_1(a+1)} + \frac{1}{k_1(b+1)} + + \frac{2}{k_2}\right)}$$

$$\Delta_m = \frac{\left(\frac{1}{k_1} + \frac{1}{k_2}\right)\left(\frac{1}{k_1(a+b+1)} + \frac{1}{k_2}\right)}{\left(\frac{1}{k_1(b+1)} + \frac{1}{k_2}\right)\left(\frac{1}{k_1(a+1)} + \frac{1}{k_2}\right)}$$

Similarly, for scheme (2), we obtain:

$$\Delta_a = \frac{\left(\frac{1}{k_1} + \frac{1}{k_2}\right)}{\left(\frac{1}{k_1} + \frac{1}{k_2(b+1)} + \frac{1}{k_1(a+1)} + \frac{1}{k_2}\right)}$$

$$\Delta_m = \frac{\left(\frac{1}{k_1} + \frac{1}{k_2}\right)\left(\frac{1}{k_1(a+1)} + \frac{1}{k_2(b+1)}\right)}{\left(\frac{1}{k_1} + \frac{1}{k_2(b+1)}\right)\left(\frac{1}{k_1(a+1)} + \frac{1}{k_2}\right)}$$

**2. Numerical Calculations.**

To gain insights the two different regulatory schemes, we evaluate the deviations from "additive" and "multiplicative" behaviours for a range of system parameters. Specifically, we select $a$ and $b$ in the range [1, 100]. We further impose a constraint of the rates of the two steps in the cascade being within ~3-fold of each other, by selecting the ratio $k_1'/k_2'$ in the range [0.3-3]. We numerically calculate $\Delta_a$ and $\Delta_m$ for $N = 1{,}000$ random sets of parameters, and visualize the results in an $xy$ scatter plot. The results show strongly "super-multiplicative" and "super-additive" behaviour ($\Delta_a$ and $\Delta_m$ both >1) for scheme (1). At the same time, we observe mostly additive and sub-additive behaviours ($\Delta_a \leq 1$ and $\Delta_m < 1$) for scheme (2). The results of these calculations are shown in Supplementary Fig. 6**B**.

**REFERENCES.**


1. D. Herschlag, F. B. Johnson, Synergism in transcriptional activation: a kinetic view. *Genes Dev* **7**, 173-179 (1993).
2. C. Scholes, A. H. DePace, A. Sanchez, Combinatorial Gene Regulation through Kinetic Control of the Transcription Cycle. *Cell Syst* **4**, 97-108 e109 (2017).
